# Two distinct mechanisms of flavoprotein spectral tuning revealed by low-temperature and time-dependent spectroscopy

**DOI:** 10.1101/2023.07.17.549366

**Authors:** Andrey Nikolaev, Elena V. Tropina, Kirill N. Boldyrev, Eugene G. Maksimov, Valentin Borshchevskiy, Alexey Mishin, Anna Yudenko, Alexander Kuzmin, Elizaveta Kuznetsova, Oleg Semenov, Alina Remeeva, Ivan Gushchin

## Abstract

Flavins such as flavin mononucleotide or flavin adenine dinucleotide are bound by diverse proteins, yet have very similar spectra when in the oxidized state. Recently, we developed new variants of flavin-binding protein CagFbFP exhibiting notable blue (Q148V) or red (I52V A85Q) shifts of fluorescence emission maxima. Here, we use time-resolved and low temperature spectroscopy to show that whereas the chromophore environment is static in Q148V, an additional protein-flavin hydrogen bond is formed upon photoexcitation in the I52V A85Q variant. Consequently, in Q148V, excitation, emission and phosphorescence spectra are shifted, whereas in I52V A85Q, excitation and low-temperature phosphorescence spectra are relatively unchanged, while emission spectrum is altered. We also determine X-ray structures of the two variants to reveal the flavin environment and complement the spectroscopy data. Our findings illustrate two distinct color tuning mechanisms of flavin-binding proteins and pave the way for engineering of new variants with improved optical properties.

**TOC GRAPHICS:** 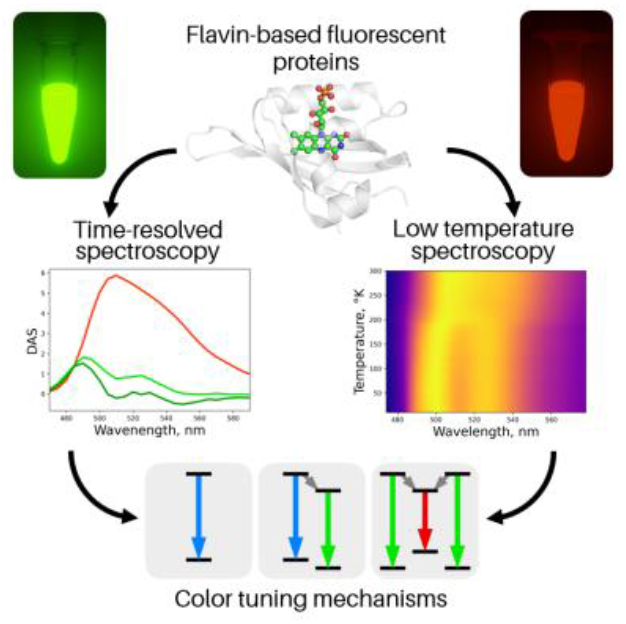

## TEXT

Flavin-binding fluorescent proteins (FbFPs) are an emerging class of fluorescent reporters for *in vivo* microscopy^1–3^. FbFPs are derived from naturally occurring Light Oxygen Voltage (LOV) domains, which act mostly as blue light sensors^4^. FbFPs typically have a brightness between approximately 3000 and 6500 M^-1^ cm^-1^ and an emission maximum in the vicinity of 497 nm^5,6^. Significant research efforts have been devoted to engineering FbFPs with altered spectra. Methods such as site-directed mutagenesis guided by QM/MM modeling, including electrostatic spectral tuning maps, or site-saturation mutagenesis have been applied by different research groups^7–11^. To date, the largest emission shifts have been obtained for a thermostable protein CagFbFP^12^, where site saturation mutagenesis allowed identification of a palette of 22 variants with emission maxima covering the range between 486 and 512 nm^10^.

Strategies for color tuning of genetically encoded fluorescent proteins can be divided into three categories: (1) altering chemical properties of chromophores, such as the protonation state or the number of conjugated bonds in π-system (relevant for the GFP family^13^); (2) altering chromophore conformation (relevant for retinylidene proteins^14^ and biliverdin-based proteins^15,16^); (3) altering polarity or polarizability of the environment of the chromophore-binding site^8^. The last approach is the only choice for stiff endogenous chromophores such as flavins.

A common strategy for color tuning of FbFPs is to add or remove charged or polar amino acids around the polar part of the isoalloxazine moiety of flavin. The effect of mutations on the emission spectrum is theoretically assessed based on the conformations of these amino acids, which are obtained from molecular dynamics simulations^7,17^ or X-ray crystallographic structures where possible^9,18^. However, redistribution of flavin charges after photoexcitation may cause changes in the conformations of surrounding polar amino acids. This phenomenon becomes especially prominent if the characteristic times of conformational changes are close to the fluorescence lifetime, and can be revealed using low-temperature and time-resolved fluorescence. These methods have been previously applied to studying the LOV domain photocycle^19^ but not yet to studying photochemical properties of FbFPs.

In this letter, we investigated the mechanisms of spectral shift of two recently discovered variants of CagFbFP displaying largest observed shifts: Q148V, which has a blue shift of 12 nm, and A85Q I52V, which has a red shift of 14 nm. A85Q introduces a polar amino acid into the chromophore binding pocket, while Q148V removes one (Fig. 1A). I52V is a compensatory mutation that does not affect the emission spectrum^10^. It is worth noting that mutation Q148V shifts both the absorbance and fluorescence spectra, whereas A85Q only shifts the fluorescence emission spectrum (Fig. 1B,C). Static amino acid environment should result in the same absorption and emission shifts. Thus, the chromophore environment in the A85Q variant might display conformational changes after flavin excitation.

**Figure 1.**
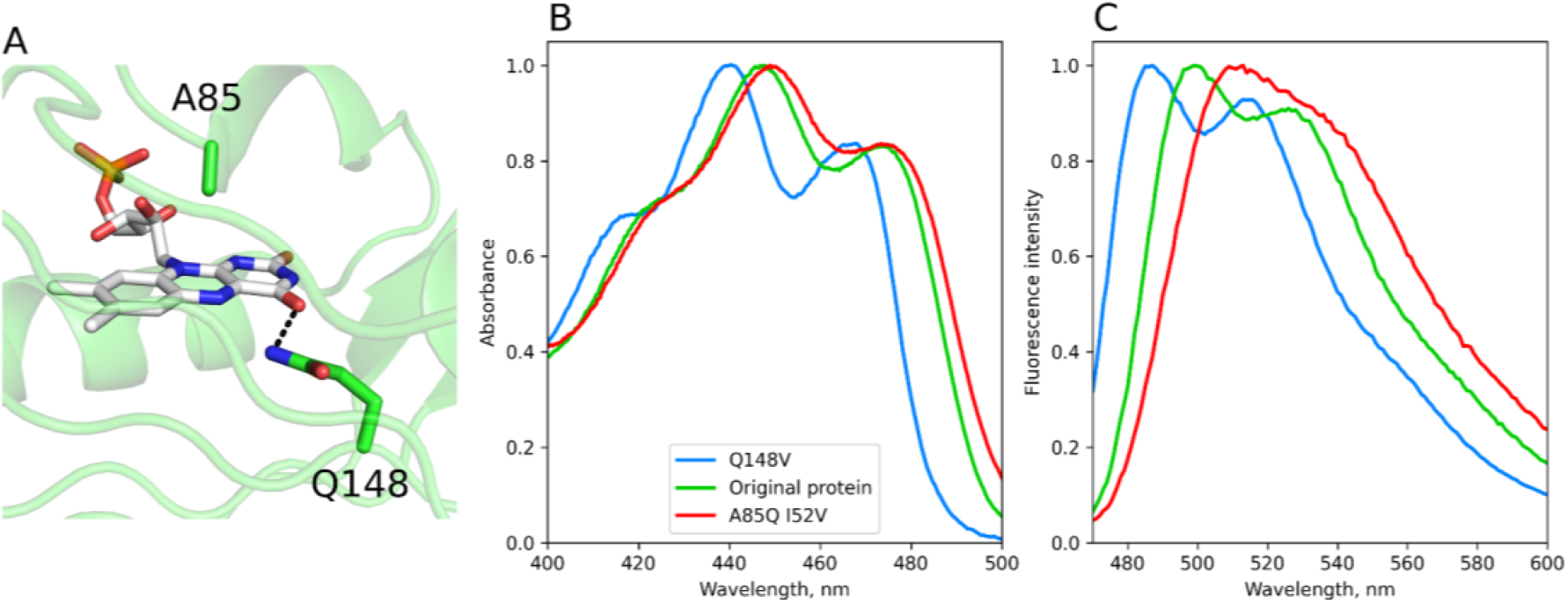
Effect of mutations on the spectroscopic properties of CagFbFP. **A**. Crystallographic structure of CagFbFP (PDB ID 6RHF^12^). **B**. Absorption spectra of Q148V and A85Q I52V variants and the original protein. While the blue shift caused by Q148V is significant, the red shift caused by A85Q is modest. **C**. Emission spectra of the three CagFbFP variants. Excitation wavelength is 450 nm. Both mutations resulted in shifts of the same magnitude. Original spectra were obtained in Ref. ^10^.

We measured time-resolved emission spectra of the original protein and the two mutants (Fig. 2). The fluorescence of the Q148V variant has a decay rate that is independent of wavelength (Fig. 2A). The decay rate of the original protein shows slight dependence on wavelength (Fig. 2E), and the decay of A85Q I52V fluorescence is significantly non-monoexponential and dependent on wavelength (Fig. 2J). We performed a global analysis of the recorded spectra and calculated the spectrum shapes the moment after the excitation and after sufficient time after the excitation (Fig. 2F,K). While the spectrum shapes of the original protein and Q148V variant are almost time-independent, the spectrum of A85Q I52V significantly shifts to longer wavelengths within 2 ns after the excitation. This time dependence can be explained by the presence of additional temporal components with smaller lifetimes (Fig. 2L).

**Figure 2.**
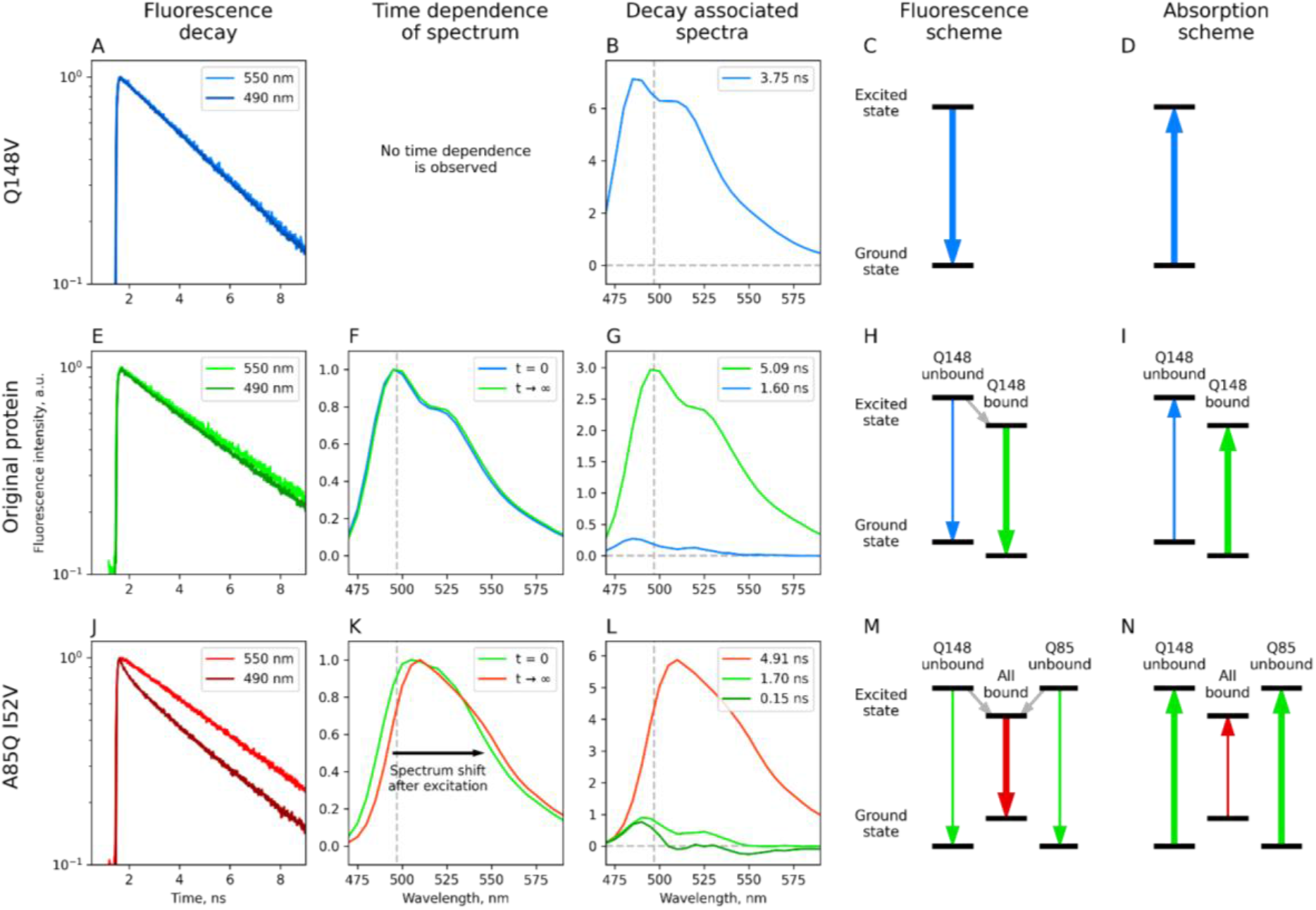
Fluorescence emission by CagFbFP and its Q148V and I52V A85Q variants and proposed energy level schemes. **A**,**E**,**J**. Fluorescence decay traces at different wavelengths. **F**,**K**. Fluorescence emission spectra immediately after excitation and after significant time. Spectra were calculated from global analysis. **B**,**G**,**L**. Decay associated spectra obtained from global analysis. One, two and three components are observed in data for Q148V, original protein, and I52V A85Q, respectively. **C**,**D**,**H**,**I**,**M**,**N**. Proposed fluorescence emission and absorption energy level schemes. Vibrational energy levels are not drawn for simplicity. The thickness of the colored arrows reflects the approximate contribution of the transition to the total fluorescence intensity or absorption. Gray arrows correspond to proposed conformational changes of the chromophore environment.

Further insights into the nature of time dependence of spectra can be obtained using low-temperature spectroscopy. Cooling the sample to temperatures of liquid helium significantly slows down or prohibits conformational changes of the amino acid environment^20^. We recorded the temperature dependence of the spectra of proteins down to 15 K (Supporting Figure 1). The spectra of the original CagFbFP and its Q148V variant display sharpening but not shift, whereas the emission spectrum of the A85Q I52V variant changes its shape below ∼200 K so that it becomes much more similar to that of CagFbFP (Fig. 3).

**Figure 3.**
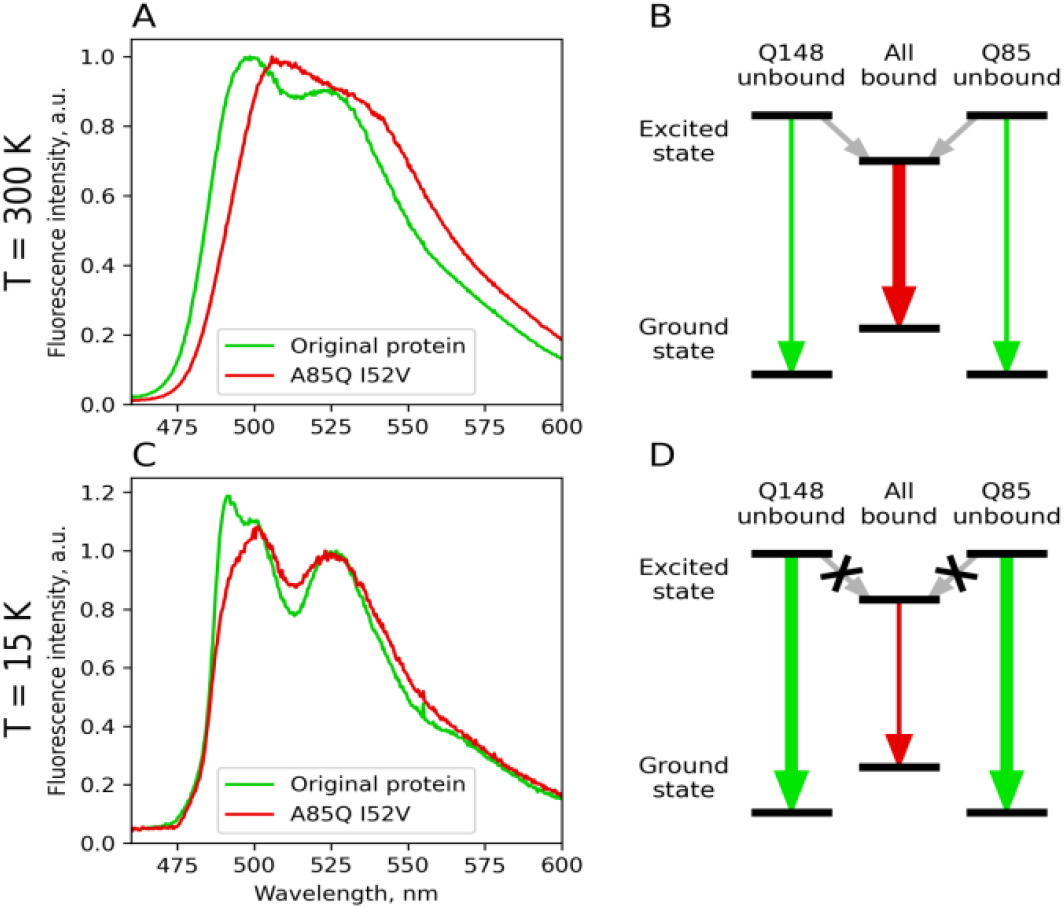
Fluorescence emission by CagFbFP I52V A85Q variant at low temperature. The red shift of the emission spectrum vanishes upon cooling. **A**,**C**. Emission spectrum of the A85Q I52V variant compared with that of the original protein at 300 K and 15 K. Upon cooling, the shape of the spectra and positions of the maxima become similar. **B**,**D**. Energy level scheme for A85Q I52V variant at 300 K and 15 K. Cooling slows down the transitions to the red-shifted conformation. The thickness of the colored arrows reflects the approximate contribution of the transition to the total fluorescence intensity.

To explain these observations, we propose energy level schemes for the original CagFbFP and its variants, which are consistent with all observed phenomena. The number of different conformations of polar amino acids in the close proximity of the chromophore would coincide with the number of temporal components. Q148V has only one temporal component (Fig. 2B), making its energy level scheme the simplest (Fig. 2C,D). Polar amino acids surrounding the chromophore (Q89, N117, and N127) remain bound to the flavin in both the ground and excited states.

The original protein (CagFbFP) has the main long-lived temporal component and a small yet noticeable shorter-lived second component with a blue-shifted spectrum (Fig. 2G). Thus, the amino acid conformation corresponding to that component is energetically unfavorable both in ground and excited states (Fig. 2H,I). The lifetime of the second component is 1.6 ns, below the fluorescence lifetime of any known FbFPs (2.5-5.5 ns). Consequently, we attribute this second component to a conformation, where Q148 is not bound to the flavin. After photoexcitation, the unbound conformation becomes even more energetically unfavorable, and its population decreases due to transition to a bound conformation. This additional decrease explains the shorter lifetime of the unbound conformation.

The fluorescence decay of the A85Q I52V variant exhibits three temporal components with linearly dependent spectra (Fig. 2L). While this linear dependence may be coincidental, it could also indicate that two of the three conformations share the same emission spectrum. The short lifetime of these two components suggests a transition from two energetically unfavorable conformations to a favored red-shifted conformation in the excited state (Fig. 2M,N). However, the variant’s almost non-shifted absorption spectrum suggests that these two conformations are energetically favored in the ground state.

Presence of the conformational transitions also explains the low temperature spectroscopy observations. Below a certain temperature, the protein does not convert to red-shifted conformation, and its spectra correspond to the spectra of the other two conformations (Fig. 3B,D). Their spectrum cannot be directly calculated from global analysis; however, based on low temperature spectra (Fig. 3C), it is similar to that of the original CagFbFP. In summary, the A85Q I52V variant (1) absorbs light in one or two non-shifted conformations; (2) converts into the red-shifted conformation, while emitting a photon before or after conversion; (3) converts back into the non-shifted conformation. Cooling below 200 K prevents conformational changes, causing the variant to absorb and emit photons in the non-shifted conformation.

During experiments with low-temperature spectroscopy, after the excitation beam was stopped, we were surprised to observe scarlet phosphorescence with a lifetime of 0.25-0.5 seconds in all studied protein samples (see Supplementary Movie 1 for CagFbFP). We measured time-resolved phosphorescence and successfully obtained decay-associated spectra with a sufficient signal-to-noise ratio by performing global analysis (Fig. 4). Q148V has blue-shifted phosphorescence compared to A85Q I52V and original protein. These shifts correspond to absorption and emission shifts at low temperature, indicating that the singlet and triplet excitation energies of flavin are similarly affected by electrostatic interactions with surrounding amino acids. Besides the main phosphorescence with an emission maximum at 600 nm, we observed weak phosphorescence with maxima around 540 nm for Q148V and the original protein, which has a smaller lifetime compared to the main maximum.

**Figure 4.**
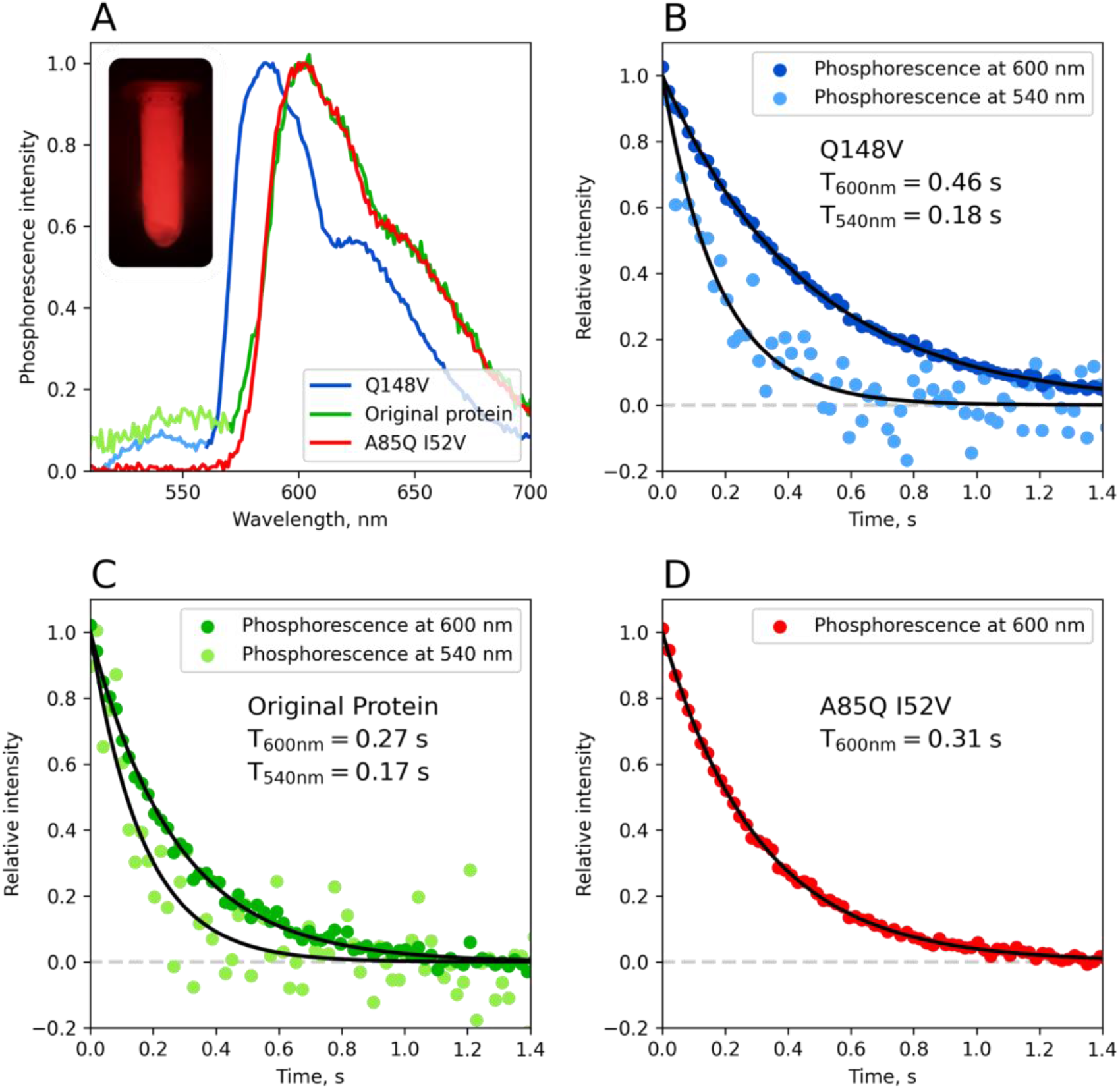
Phosphorescence emission spectra and decay curves of CagFbFP variants at 50 K. **A**. Decay associated spectra for the original protein and its variants obtained from monoexponential global analysis. *Inset:* Photograph of phosphorescence of CagFbFP in a 2 ml tube after switching off the excitation light at 77 K. **B-D**. Phosphorescence decay curves for Q148V, original, and I52V A85Q variants at 600 and 540 nm (where present), respectively.

To gain further insight into the color tuning mechanisms at the atomic level, we determined crystallographic structures of Q148V and I52V A85Q variants (Fig. 5). The overall structures of mutants are identical to those of the original variant CagFbFP^12^ in all regards except the side chains of mutated amino acids; root mean square deviations of backbone atoms are 0.35 and 0.4 Å, correspondingly. In the Q148V variant, the only difference is the mutation, and V148 is observed in different rotameric states (Fig. 5B). In the I52V A85Q variant, all three active site residues assume multiple conformations. In the first conformation (Fig. 5D), the Q85 amide group is co-planar with the flavin, and the N_ε2_ of Q148 makes hydrogen bonds both with O_ε1_ of Q85 and O4 of the flavin. In the second conformation (Fig. 5E), Q148 is slightly displaced away from Q85, whereas Q85 amide is rotated so that N_ε2_ of Q85 is hydrogen-bonded to flavin’s N5, and N_ε2_ of Q148 is hydrogen-bonded to flavin’s O4. The correspondence between the rotameric states of V52 and the conformations of Q85 and Q148 is not clear.

**Figure 5.**
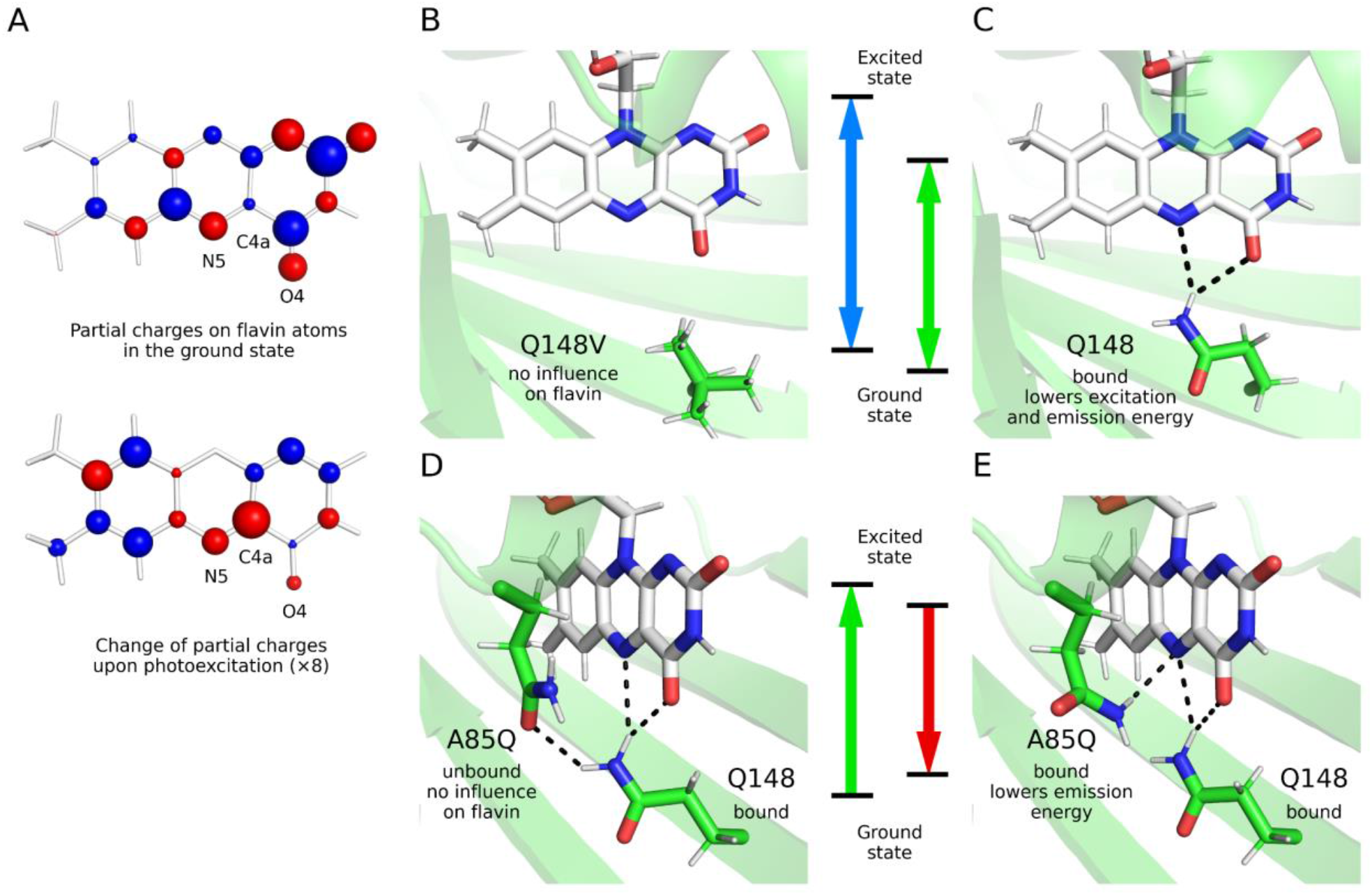
Structural insights into the mechanisms of color tuning in CagFbFP mutants. **A**. Partial charges on FMN isoalloxazine moiety atoms (top) and change of these partial charges upon photoexcitation (bottom), with red and blue representing negative and positive charges, respectively, and radius of the sphere proportional to the square root of the respective values. Hydrogen charges were added to the charges of the respective heavy atoms. **B**. Crystallographic structure of the Q148V variant. No protein atoms form hydrogen bonds with N5 and O4 atoms of the flavin. **C**. Crystallographic structure of the original protein (PDB ID 6RHF^12^). Electron redistribution to the O4, C4a and N5 atoms of the flavin upon excitation is favored in the original protein due to proximity of the partially positively charged Q148 side chain amide group. **D**,**E**. Two alternative conformations of Q148 and Q85 side chains observed in crystallographic structure of the I52V A85Q variant. Charge redistribution upon excitation favors reorientation of Q85 side chain and formation of a hydrogen bond between A85Q amide group and N5 atom of the flavin.

To understand the mechanism of spectral shifts at the atomic level, we calculated the difference of the flavin’s partial atomic charges between the ground and excited state (Fig. 5A). In accordance with previous studies^5,8^, we conclude that the blue shift of the Q148V variant results from the loss of the hydrogen bond between Q148 and flavin. Flavin atoms close to the amide group of Q148 gain additional negative charge after photoexcitation. Presence of partially positively charged hydrogen atom of the amide group of Q148 nearby favors this charge redistribution, which leads to lowering of the excitation and emission energy. In the original CagFbFP, photoexcitation favors conversion of the small population, where Q148 is not bound to the flavin, to the conformation where Q148 is bound to the flavin, which is accompanied by shortening of the fluorescence lifetime of the blue-shifted unbound state and displayed as time-dependent shift of the fluorescence emission maximum towards 498 nm.

For the I52V A85Q variant, the first conformation is likely to be spectrally similar to that of the original protein CagFbFP, since Q148 is bound to the flavin and Q85 is not. The second conformation should be red-shifted, since Q85 forms a hydrogen bond with flavin’s N5, favoring electron redistribution towards N5, which happens upon photoexcitation. Finally, it is possible that in solution, similarly to original CagFbFP, a third conformation is observed, where Q148 is not bound to the flavin. We reconcile the structural and spectroscopic data by suggesting that in the darkness the protein mostly assumes the conformations where either Q85 or Q148 is not bound to the flavin, probably due to steric conflicts. Upon illumination, the protein converts to the conformation where both Q85 and Q148 are hydrogen-bonded to the flavin. Because the transition relies on flipping of the glutamine side chains, it is inhibited by temperatures below the glass transition of proteins^20^.

To summarize, we studied two variants of the CagFbFP protein and found that they have distinct mechanisms of the emission spectrum shifts. For Q148V, the chromophore environment is static, and the emission spectrum is shifted similarly to the excitation spectrum. For I52V A85Q, photoexcitation favors formation of a second hydrogen bond between one of the amide moieties of adjacent glutamines and the flavin, resulting in time-dependent red shift of the emission spectrum. Characteristic time of the transition from the ground state protein conformation to the excited state protein conformation is shorter than the fluorescence lifetime for CagFbFP, and thus the conformational change has a significant impact on the emission spectrum.

Presented findings provide a comprehensive explanation of the photophysics and photochemistry underlying the newly developed palette of FbFP proteins^10^ and highlight low-temperature and time-resolved spectroscopy as fruitful approaches for studying FbFP color tuning. We hope that this work will pave the way for engineering of new flavin-binding protein variants with improved optical properties, not limited to fluorescent proteins, but also including optogenetic tools^2^, singlet oxygen generators^21^ and photoenzymes^22,23^.

## Methods

### Protein expression and purification

CagFbFP and Q148V and A85Q I52V mutants were expressed and purified as described previously^10,12^. Proteins were transferred to a buffer containing 100 mM NaCl, 20 mM Tris-HCl pH 8.0 for low-temperature spectroscopy and to a buffer containing 300 mM NaCl, 50 mM Tris-HCl pH 8.0 for time-resolved resolved fluorescence spectroscopy by dialysis.

### Low-temperature spectroscopy

Samples were cooled to 10-15 K using PT403 closed cycle helium cooled cryostat (CryoMech, USA). Fluorescence excitation was performed with an ultraviolet diode with an emission range of 360-370 nm and power density of 50 mW/cm^2^. Spectra were recorded using the HR-series spectrometer (Ocean Insight, USA) with a spectral resolution of 1.75 nm.

The decay of phosphorescence was recorded using the same experimental setup at 50 K. The sample was illuminated with a diode for a few seconds, after which the diode was momentarily switched off while the emission spectra were recorded every 20 milliseconds. The spectra related to the phosphorescence decay were selected based on the absence of fluorescence at 500 nm.

### Time-resolved fluorescence spectroscopy

A set of fluorescence decay kinetics was measured in a time-correlated single photon (TCSPC) mode using a detector with an ultra-low dark count rate (HPM100-07C, Becker&Hickl, Germany) coupled to a ML-44 monochromator (Solar LS, Belarus) used for tuning of the detection wavelength from 470 to 600 nm in 5 nm steps. Fluorescence was excited at 440 nm (repetition rate 80 MHz, pulse width 150 fs, average optical power 1 mW) using the second harmonics of a femtosecond optical parametric oscillator (TOPOL-1050-C, Avesta Project LTD, Russia) pumped by a femtosecond Yb-laser (TEMA-150, Avesta Project LTD, Russia). The emission signal was collected perpendicular to the excitation beam. To avoid the detection of scattered laser light, the edgepass filter (long-pass 450 nm Thorlabs, USA) was used. The temperature of the samples was stabilized at 25 °C by a thermostatic cuvette holder Qpod 2e with a magnetic stirrer (Quantum Northwest, USA).

### Global analysis of time-resolved spectra

Time resolved fluorescence data was fitted using following functions:

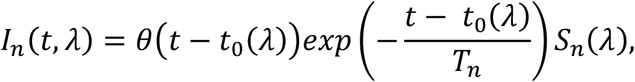

where *T*_*n*_ is life-time of temporal component, *S*_*n*_(*λ*) is decay associated spectra, *t*_0_(*λ*) is time of excitation pulse with *λ*-dependent dispersion correction, *θ*(*t*) is Heaviside step function and n is the index of the temporal component. To account for the small but noticeable residual fluorescence from the previous pulse, the following correction was made for total intensity:

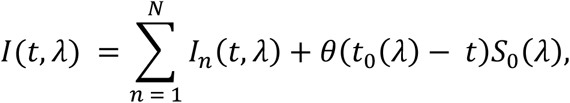

where *S*_0_(*λ*) is spectra of residual fluorescence and *I*(*t, λ*) is total intensity that was further convoluted with instrument response function (IRF) that was fitted as follows:

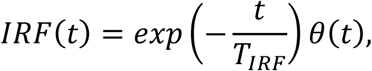

where *T*_*IRF*_ was a free parameter in the fitting procedure and was between 41 and 43 ps depending on measurement. We found that this form of approximate IRF best matches the experimental IRF measured on water sample.

Time resolved phosphorescence data was fitted using following function:

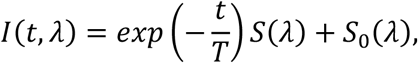

where *I*(*t, λ*) is total intensity, *S*(*λ*) is decay associated spectrum, *S*_0_(*λ*) is baseline spectrum and T is phosphorescence lifetime. Due to the low signal to noise ratio further complications including second temporal component or accounting for IRF leads to overfitting. The main body of spectra with a maximum around 600 nm and the low intensity peak around 540 nm were fitted separately.

### X-ray crystallography

Proteins were crystallized by sitting drop vapor diffusion approach using the NT8 robotic system (Formulatrix, USA). The drops contained 200 nL concentrated protein solution and 200 nL reservoir solution. Crystallization plates were stored at 20 °C. The best crystals of the Q148V variant were obtained using the following solution as a precipitant: 0.2 M Magnesium chloride hexahydrate, 0.1 M TRIS hydrochloride pH 8.5, 30% w/v Polyethylene glycol 4,000. The best crystals of the I52V A85Q variant were obtained using the following solution as a precipitant: 0.2 M Calcium acetate hydrate, 0.1 M Sodium cacodylate trihydrate pH 6.5, 18% w/v Polyethylene glycol 8,000. Elongated plate-like crystals reached the final size of ∼100-300 μm within a few weeks. The crystals were harvested using micromounts, flash-cooled and stored in liquid nitrogen. Diffraction data were collected at 100 K at the Elettra synchrotron beamline XRD2^24^ equipped with a PILATUS2 6M detector (Dectris, Switzerland). Diffraction images were processed using XDS^25^. AIMLESS^26^ was used to merge and scale the data. The data collection and processing statistics are reported in Supporting Table 1. The structures were solved using molecular replacement with MOLREP^27^ and CagFbFP structure (PDB ID 6RHF^12^) as a search model. The resulting models were refined manually using Coot^28^ and REFMAC5^29^ and deposited into Protein Data Bank under accession codes 8PKY and 8PM1.

### QM/MM modelling

Initial coordinates for modeling were taken from chain A of crystallographic structure of CagFbFP (PDB IF 6RHF^12^). Protonated structure of monomeric protein was prepared using CHARMM-GUI PDB Reader & Manipulator tool^30–32^. Protonation states of titratable residues were assigned to pH = 8 based on pK_a_. The phosphate group of FMN ligand was deprotonated to -2*e* charge. The resulting charge of CagFbFP and FMN complex was -2*e*. The system was neutralized by adding 2 Na^+^ ions. The protein was solvated in a cubic water box with a water layer of 12 Å around the protein using CHARMM-GUI Solution Builder^33,34^, while keeping the water molecules present in the crystallographic structure. For initial MD preparation of structure Amber14SB^35^ parameters and General Amber Force Field 2 (GAFF2)^36,37^ parameters were used for protein and FMN, respectively. Molecular dynamics simulations were conducted using OpenMM 7.6^38^. System was energy minimized using the steepest descent method. Then, 100 ps of equilibration were performed in order to properly equilibrate crystallographic structure with generated solvent. After that system was energy minimized again for further use in QM/MM modeling. Langevin integrator with a time step of 3 fs, friction coefficient of 1 ps and reference temperature of 300K was used. Pressure was kept at 1 bar using Monte-Carlo barostat^39^, which was applied every 25 steps. Cutoff of 1 nm were implemented for calculating Lennard-Jones interaction and particle mesh Ewald method^40^ was used for long-range electrostatic interactions with error tolerance of 0.0005. Length of bonds containing hydrogen were constrained during the simulation.

QM/MM modeling was conducted using ORCA 5.0.3^41–43^. Additive QM/MM scheme with electrostatic embedding was applied. Prepared structure was divided into two subsystems: QM subsystem containing lumiflavin and MM subsystem containing ribose-5′-phosphate tail, protein, water and ions. For the QM part of the QM/MM calculation, B3LYP^44–47^ density functional for ground state and time-dependent^48,49^ B3LYP for excited state with ma-def2-SVP^50,51^ basis set were used. Bonded interaction between subsystems was managed using a link hydrogen atom and charge-shift scheme. QM/MM-optimized geometry were obtained for both ground and first singlet excited state. All MM atoms farther than 8 angstroms from lumiflavin were fixed during optimization. Partial atomic charges of lumiflavin were calculated using the CHELPG approach^52RACTp^ for both states.

## Supporting Information

The following files are available free of charge.

Supporting Information file that includes low-temperature fluorescence emission data for all variants and X-ray crystallography statistics (PDF)

Supporting Movie 1 showing CagFbFP phosphorescence (MPEG)

## Notes

The authors declare no competing financial interests.

## ACKNOWLEDGMENT

We gratefully acknowledge Elettra synchrotron and XRD2 beamline for providing beamtime and support under proposal 20225306. Study design, sample preparation and data processing were supported by the Russian Science Foundation (project 21-64-00018). Low-temperature spectroscopy was supported by the Russian Science Foundation (project 19-72-10132P awarded to K.N.B. and E.V.T.). Protein crystallization and data collection were supported by the Russian Ministry of Science and Higher Education (grant no. 075-15-2021-1354).

## Notes

### Competing Interest Statement

The authors have declared no competing interest.

