## Supporting Information for "Two distinct mechanisms of flavoprotein spectral tuning revealed by low-temperature and time-dependent spectroscopy"

Supporting Figure 1

Supporting Table 1

Supporting Movie 1: separate file

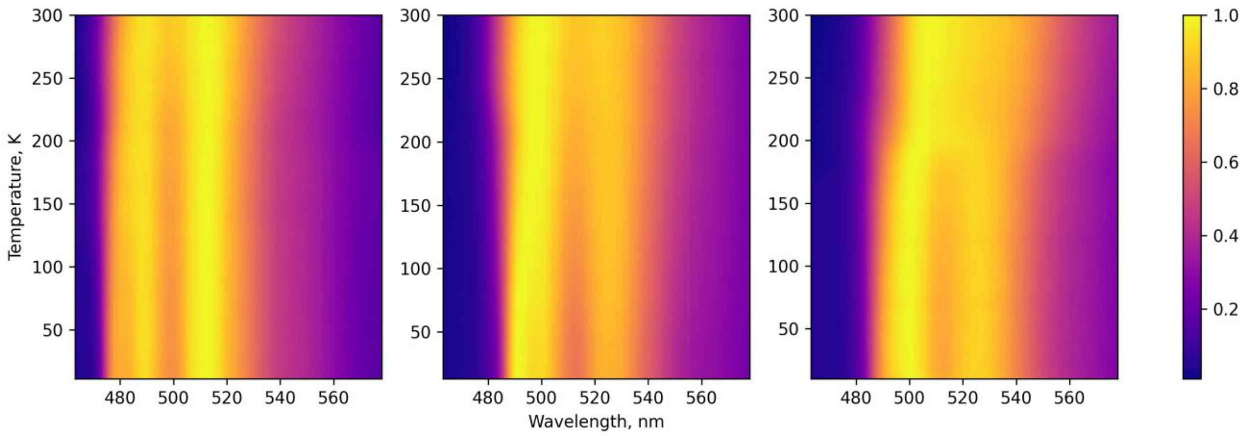


**Supporting Figure 1.** Dependence of the normalized fluorescence emission intensity depending on temperature for Q148V (left), original (center), and I52V A85Q (right) variants of CagFbFP. For the latter variant, an abrupt change in the fluorescence emission spectrum is observed when the temperature is lowered below 200 K.

**Supporting Table 1.** Crystallographic data collection and refinement statistics.

| **Variant**<br>**PDB ID** | **Q148V**<br>**8PKY** | **I52V A85Q**<br>**8PM1** |
|---|---|---|
| **Data collection** | | |
| Space group | $P2_12_12$ | $P2_12_12$ |
| Cell dimensions | | |
| *a*, *b*, *c* (Å) | 53.80, 109.39, 39.02 | 53.82, 110.43, 39.31 |
| *α*, *β*, *γ* (°) | 90, 90, 90 | 90, 90, 90 |
| Resolution (Å) | 54.75-1.40 (1.42-1.40) | 55.22-1.50 (1.53-1.50) |
| $<I/\sigma I>$ | 18.3 (1.0) | 8.5 (0.7) |
| *CC1/2 (%)* | 100.0 (41.2) | 99.8 (44.7) |
| Completeness (%) | 98.0 (96.1) | 98.9 (98.0) |
| Multiplicity | 14.1 (14.2) | 14.0 (14.0) |
| Unique reflections | 45300 (2160) | 37959 (1833) |
| **Refinement** | | |
| Resolution (Å) | 54.75-1.40 | 55.22-1.50 |
| No. reflections | 43020 | 36039 |
| $R_{work}/R_{free}$ (%) | 14.2/18.5 | 19.7/22.8 |
| No. atoms | | |
| Protein | 1707 | 1730 |
| FMN | 62 | 62 |
| Water and others | 285 | 226 |
| Average *B* factors (Å$^2$) | | |
| Protein | 21.3 | 18.3 |
| FMN | 17.5 | 13.4 |
| Water and others | 37.1 | 28.7 |
| R.m.s. deviations | | |
| Protein bond lengths (Å) | 0.003 | 0.009 |
| Protein bond angles (°) | 0.9 | 1.6 |
| Ramachandran analysis | | |
| Favored (%) | 99 | 99 |
| Outliers (%) | 0 | 0 |

* The data for the lowest and highest resolution shells are shown in parentheses. R.m.s.: root mean square.
